# Excitability modulations of somatosensory perception do not depend on feedforward neuronal population spikes

**DOI:** 10.1101/2024.07.15.603430

**Authors:** T. Stephani, A. Villringer, V. V. Nikulin

## Abstract

Neural states shape perception at earliest cortical processing levels. Previous work in humans showed a relationship between initial cortical excitation, as indicated by the N20 component of the somatosensory evoked potential (SEP), pre-stimulus alpha oscillations, and the perceived intensity in a somatosensory discrimination paradigm. Here we address the follow-up question whether these excitability dynamics reflect changes in feedforward or feedback signals. We leveraged high-frequency oscillations (HFO) as a metric for neuronal population spiking activity of the first excitatory feedforward volley in the somatosensory cortex, non-invasively extracted from electroencephalography (EEG) data of 32 male human participants. Using Bayesian statistics, we found evidence against the involvement of HFO in moment-to-moment variability of perceived stimulus intensity, in contrast to previously observed pre-stimulus alpha and N20 effects. Given that the N20 component presumably reflects backpropagating membrane potentials towards the apical dendrites, we argue that top-down feedback processes (e.g., related to alpha oscillations) may thus rely on modulations at distal dendritic sites of involved pyramidal cells rather than on output firing changes at their basal compartments.

## Introduction

The perception and neural encoding of even basic stimulus features, such as their intensity, is variable and depends on instantaneous states of the neural system (Arieli et al., 1996; Sadaghiani et al., 2010; McCormick et al., 2020). In line with the recent proposal that this variability is related to dynamics of excitability levels (Samaha et al., 2020), we found earliest cortical excitation, as measured by the N20 potential in the human somatosensory system, to depend on pre-stimulus alpha oscillations and to affect the subjectively perceived stimulus intensity (Stephani et al., 2021). Yet, whether these excitability dynamics reflect a modification of feedforward processing or are implemented via top-down feedback signals remains elusive.

The N20 component of the somatosensory evoked potential (SEP) reflects excitatory processes of the first thalamo-cortical volley arriving in the primary somatosensory cortex (S1) in Brodmann area 3b (BA 3b; Allison et al., 1991; Nicholson Peterson et al., 1995; Wikström et al., 1996; Bruyns-Haylett et al., 2017). This initial thalamo-cortical excitation starts in middle layers of S1 (i.e., layer 4), leading to a depolarization of basal dendrites near the soma of pyramidal cells (in layer 2/3 and potentially also in layer 5; Kulics and Cauller, 1986). Importantly, the anterior-positive dipole topography suggests that the N20 potential spreads from the basal compartments of involved pyramidal cells (where sensory inputs arrive) not only to the soma where axonal action potentials are generated but also along the membranes towards the apical dendrite of the pyramidal cell (Fig. 1; Wikström et al., 1996). This notion of a backpropagating membrane depolarization underlying the generation of N20 has been confirmed by invasive recordings in non-human animals (Ikeda et al., 2005), as well as by recent biophysical modeling work (Thorpe et al., 2024). Thus, the N20 most likely spans several compartments of the involved pyramidal cells.

**Fig. 1.**
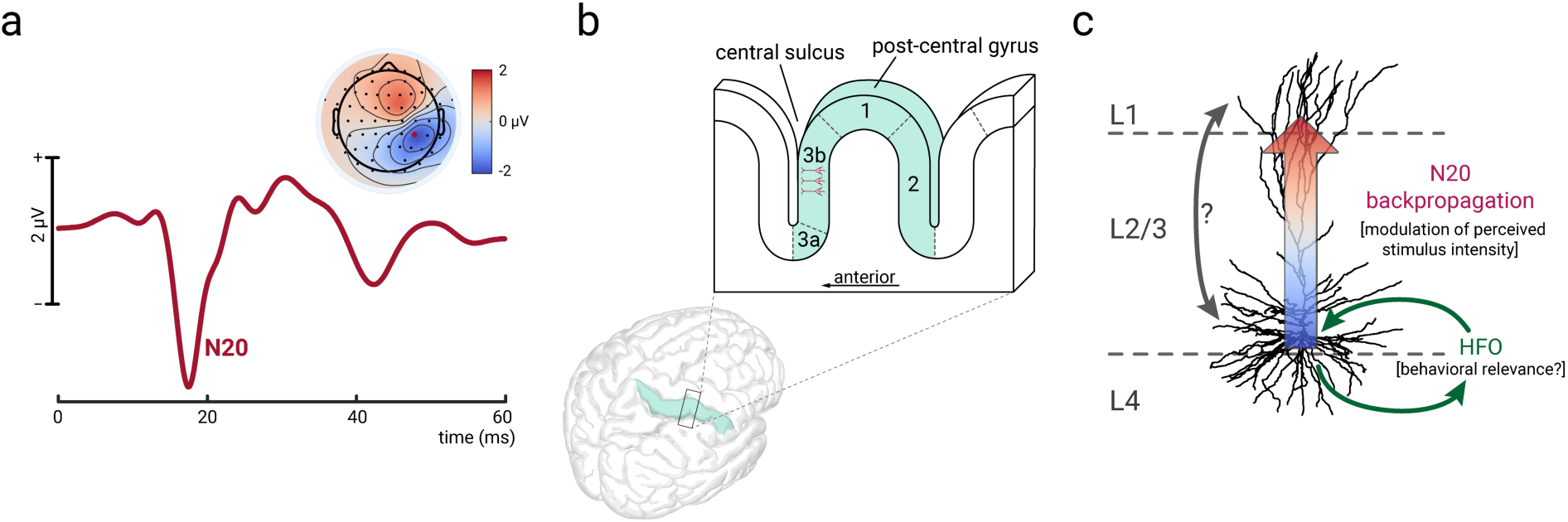
The N20 component of the somatosensory evoked potential (SEP) and its assumed generation. a) N20 component measured on scalp level with electroencephalography (EEG). SEP at electrode CP4, averaged over 1000 trials (exemplary participant). b) Location of pyramidal cells in Brodmann area 3b in the anterior wall of the post-central gyrus, which are believed to be the sources of the N20 potential. c) The N20 component emerges due to membrane depolarizations that propagate back from the basal compartments towards the apical dendrites. High-frequency oscillations (HFO) emerge in middle layers, representing neuronal population spikes that are most likely involved in feedforward inhibition circuits affecting the basal compartments (Ozaki and Hashimoto, 2011; Hu et al., 2023).

Intriguingly, distinct functional roles have been posited for different pyramidal cell compartments (Larkum, 2013): While bottom-up sensory input arrives at basal dendrites near the cell soma, the apical dendrites in superficial layers receive top-down signals from corticocortical and thalamocortical feedback loops (i.e., contextual information). In a “coupling zone” in the proximal apical dendrite, the integration of these two information streams takes place (Aru et al., 2020), utilizing the backpropagation of membrane potentials from basal towards apical compartments (Larkum et al., 1999; Larkum et al., 2004). Given that this enables an interaction between bottom-up sensory input and top-down modulatory signals (Phillips et al., 2015; Bachmann et al., 2020; Aru et al., 2020; Marvan et al., 2021), the question arises whether the somatosensory excitability fluctuation that we recently found in N20 responses (Stephani et al., 2021) took place via a similar mechanism of top-down modulatory states in superficial layers or rather reflected changes of feedforward output firing in middle and/or deep layers.

While EEG cannot resolve layer-specific cortical activity directly, high-frequency oscillations (HFO; >500 Hz) in response to median nerve stimulation allow to study neuronal population spiking activity generated in regions of the basal pyramidal cell compartments in S1 (Shimazu et al., 2000; Curio, 2000; Baker et al., 2003). Based on exact timing (at or just after the N20 peak; Ritter et al., 2008) and tangential dipole characteristics (Gobbelé et al., 2008), a subclass of these HFO can be distinguished that likely originate from the same pyramidal cells as the N20 potential (also shown by invasive recordings in monkeys and piglets; Shimazu et al., 2000; Ikeda et al., 2005).

Re-analyzing a previous EEG dataset of a somatosensory intensity discrimination task (Stephani et al., 2021), we found evidence *against* the involvement of HFO in the modulation of perceived stimulus intensity – in contrast to pre-stimulus alpha and N20 amplitudes – suggesting that behaviorally relevant modulations of cortical excitability are not likely to be manifested in neuronal population spikes that occur around the basal compartments of S1 pyramidal cells, but rather take place at apical sites of pyramidal cells at this initial processing stage in S1.

## Materials and Methods

This work builds on the dataset and part of the analyses of Stephani et al., 2021, published in *eLife* [link to *Materials and methods*: https://elifesciences.org/articles/67838#s4]. EEG pre-processing and signal extraction of the N20 response of the somatosensory evoked potential (SEP), as well as of pre-stimulus alpha band activity, were identical as in our previous study. Specific additional pre-processing steps were taken to extract high-frequency oscillations (HFO), and Bayesian statistics were employed to evaluate the evidence for the existence and non-existence of the respective brain-behavior relationships.

### Participants

32 male human participants (mean age = 27.0 years, *SD* = 5.0) were recruited from the database of the Max Planck Institute for Human Cognitive and Brain Sciences, Leipzig, Germany. All participants were right-handed (lateralization score, *M* = +93.1, *SD* = 11.6; assessed using the Edinburgh Handedness Inventory, Oldfield, 1971) and without any reported neurological or psychiatric disease. All participants gave informed consent and were reimbursed monetarily. The study was approved by the local ethics committee (Ethical Committee at the Medical Faculty of Leipzig University, 04006 Leipzig, Germany).

### Stimuli

Somatosensory stimuli were delivered via electrical stimulation of the median nerve using a non-invasive bipolar stimulation electrode placed on the left wrist (cathode proximal). For the stimulation, square electrical pulses with a duration of 200 µs were generated with two DS-7 constant-current stimulators (Digitimer, Hertfordshire, United Kingdom). Two stimulus intensities were used, in the following referred to as *weak* and *strong* stimulus. The weak stimulus was set at 1.2 times the motor threshold, producing a clearly visible thumb twitch with each stimulation. The motor threshold was determined as the lowest intensity that caused a visible thumb twitch, identified through a staircase procedure. The strong stimulus was calibrated during pre-experiment training blocks, set just above the participant’s *just-noticeable difference*, corresponding to a discrimination sensitivity of about *d’* = 1.5, based on Signal Detection Theory (SDT; Green and Swets, 1966). Although both stimuli were clearly perceptible, they were only minimally distinguishable, with average intensities of 6.60 mA (*SD* = 1.62) and 7.93 mA (*SD* = 2.06), for the weak and strong stimulus, respectively.

### Experimental Procedure

During the experiment, participants were comfortably seated in a chair with their hands extended in front of them, palms facing upward on a pillow. The left hand and wrist, where the stimulation electrodes were attached, were covered by a paper box to prevent the participants from visually assessing the stimulus intensity by the extent of thumb twitches caused by the stimulation. Weak and strong stimuli were presented with equal probability in a continuous, pseudo-randomized sequence with inter-stimulus intervals (ISI) ranging from 1463 to 1563 ms, drawn randomly from a uniform distribution (ISI_average_ = 1513 ms). A total of 1000 stimuli were administered, divided into five blocks of 200 stimuli each with short breaks between blocks. After each stimulus, participants had to indicate whether they felt the weak or strong stimulus by pressing a button with their right index or middle finger as quickly as possible. The button assignment for weak and strong stimulus was counterbalanced across participants. Additionally, each sequence began with a weak stimulus to serve as a reference point for the intensity judgments, and participants were informed about this. While performing the discrimination task, participants were instructed to fix their gaze on a cross on a computer screen in front of them.

Before the main experiment, training blocks of 15 stimuli each were conducted to familiarize the participants with the task and to individually adjust the intensity of the strong stimulus to achieve a discrimination sensitivity of approximately *d’*=1.5. On average, participants completed 10.5 training blocks (SD=5.8). During these training sessions, participants received visual feedback on the accuracy of their responses, but no performance feedback was provided during the experimental blocks.

### Data acquisition

EEG data were collected from 60 Ag/AgCl electrodes at a sampling rate of 5000 Hz using an 80-channel EEG system (NeurOne Tesla, Bittium, Oulu, Finland). A built-in band-pass filter in the frequency range from 0.16 to 1250 Hz was applied. The electrodes were arranged in an elastic cap (EasyCap, Herrsching, Germany) following the international 10-10 system at positions FP1, FPz, FP2, AF7, AF3, AFz, AF4, AF8, F7, F5, F3, F1, Fz, F2, F4, F6, F8, FT9, FT7, FT8, FT10, FC5, FC3, FC1, FC2, FC4, FC6, C5, C3, C1, Cz, C2, C4, C6, CP5, CP3, CP1, CPz, CP2, CP4, CP6, T7, T8, TP7, TP8, P7, P5, P3, P1, Pz, P2, P4, P6, P8, PO7, PO3, PO4, PO8, O1, and O2, with FCz as the reference and POz as the ground. Additionally, four electrodes were positioned around the eyes—at the outer canthus and the infraorbital ridge of each eye—to record the electrooculogram (EOG). Electrode impedance was kept below 10 kΩ. Furthermore, the compound nerve action potential (CNAP) of the median nerve in the left upper arm and the compound muscle action potential (CMAP) of the left M. abductor pollicis brevis were measured. These data are reported elsewhere (Stephani et al., 2021).

### EEG preprocessing and canonical correlation analysis for single-trial extraction

Stimulation artifacts were interpolated between –2 to 4 ms relative to stimulus onset using Piecewise Cubic Hermite Interpolating Polynomials (MATLAB function *pchip*). For the extraction of the conventional SEP, including the N20 component, the EEG data were band-pass filtered between 30 and 200 Hz, applying a 4^th^ order Butterworth filter in both forward and backward direction to prevent phase shifts (MATLAB function *filtfilt*). As described in Stephani et al. (2021), this filter was used to specifically extract the N20-P35 complex of the SEP, which arises at frequencies above 35 Hz, while excluding later SEP components that were not of interest. Additionally, this filter effectively acted as a baseline correction, removing slow trends in the data with an attenuation of 30 dB at 14 Hz, ensuring that SEP fluctuations were not spuriously contaminated by slower frequencies (e.g., alpha band activity). Data segments affected by muscle or non-biological artifacts were visually identified and removed. The data were then re-referenced to an average reference and eye artefacts were removed using independent component analysis (Infomax ICA). The ICA weights were calculated on the data band-pass filtered between 1 and 45 Hz using a 4^th^ order Butterworth filter applied forward and backward direction. For SEP analysis, the data were segmented into epochs from –100 to 600 ms relative to stimulus onset, resulting in an average of 995 trials per participant. EEG pre-processing was performed using EEGLAB (Delorme and Makeig, 2004), and custom MATLAB scripts (The MathWorks Inc., Natick, Massachusetts).

Single-trial SEPs were extracted using Canonical Correlation Analysis (CCA), following the approach of Waterstraat et al. (2015), and applied similarly as in Stephani et al. (2020) for a comparable dataset.

CCA finds the spatial filters ***w_x_*** and ***w_y_***_”_ for multi-channel signals ***X*** and ***Y*** by solving the following optimization problem for maximizing the correlation:

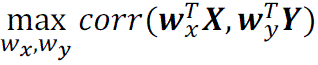

where ***X*** is a multi-channel signal constructed from concatenating all the epochs of a subject’s recording, i.e. ***X*** = [***x***_1_, ***x***_2_, …, ***x_x_***] with ***x_i_*** ∈ ℝ^channel×.time^ being the multi-channel signal of a single trial and *N* the total number of trials. Additionally, 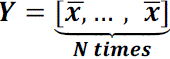 with 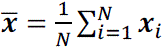 denoting the grand average of all trials. The spatial filter ***w_x_*** represents a vector of weights for mixing the channels of each single trial (i.e. 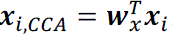) that allows to recover their underlying SEP. As we focused on the early portion of the SEP, the signal matrices ***X*** and ***Y*** were constructed using post-stimulus segments from 5 to 80 ms. Nevertheless, the extracted CCA spatial filter was applied to the full-length epochs from –100 to 600 ms. Applying the CCA spatial filter ***w_x_*** to the single trialś signals results in a set of CCA components, 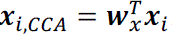. Multiplying the spatial filters ***w_x_*** by the covariance matrix of ***X***, i.e. *cov*(***X***)***w_x_***, leads to the spatial patterns of each CCA component (Haufe et al., 2014). For further analysis, we selected the CCA components whose spatial patterns was consistent with the presence of a tangential dipole over the central sulcus, as is typical for the N20-P35 complex, referred to in the following as *tangential CCA components*. For all participants, such a tangential CCA component was present among the first two CCA components with the maximum canonical correlation coefficients. Accounting for the insensitivity of the CCA solutions to the polarity of the signal, the resulting tangential CCA components were standardized by multiplying the spatial filter by a sign factor so that the N20 potential always appeared as a negative peak in the SEP.

Identically as in the previous work (Stephani et al., 2021), we extracted N20 peak amplitudes as the most negative value in single-trial SEPs of the tangential CCA components ±2 ms around the latency of the N20 in the within-subject average SEP. Pre-stimulus alpha band amplitudes were extracted in a time window of –200 an –10 ms relative to stimulus onset, after segmenting the data from –500 to –5 ms and applying a band-pass filter between 8 and 13 Hz (4^th^ order Butterworth filter applied forwards and backwards). Filter-related edge artifacts were minimized by mirroring the data segments before filtering to both sides. Next, the spatial filter of the tangential CCA component was applied to the pre-stimulus alpha data to extract comparable neuronal sources. Hilbert transform was applied to the real-valued signal of the pre-stimulus alpha oscillations, allowing to estimate the amplitude envelope by taking the absolute values of the analytic signal. One pre-stimulus alpha metric was obtained for every trial, by averaging the alpha envelope in the pre-stimulus time window between –200 and –10 ms. Log-transformation was applied in order to approximate a normal distribution for subsequent statistical analyses.

### Pre-processing and signal extraction of HFO

General EEG pre-processing for the extraction of high-frequency oscillations (HFO) was similar as in the above described N20 analysis yet deviating regarding the EEG frequency range of interest: The EEG data were band-pass filtered between 550 and 850 Hz with a sixth-order Butterworth filter (applied forwards and backwards to prevent phase shift). This filter was tuned to HFO activity centered around 700 Hz with a range of ±150 Hz (as was indicated to be appropriate by prior time-frequency analyses). The same data segments and the same ICA components were removed as in the previous analyses, leading to the same amount of trials of HFO activity as in our original work.

Again, we used canonical correlation analysis (CCA) to find spatial filters that optimized the signal-to-noise ratio of single-trial evoked activity – a method that was developed for HFO specifically (Fedele et al., 2013; Waterstraat et al., 2015). Deviating from our previous analysis of conventional SEP (i.e., N20 component), we here trained the CCA weights in time windows from 10 to 30 ms (but, again, applied the resulting spatial filters to the whole data epochs from –100 to 600 ms). The resulting CCA components were visually inspected regarding their stimulus-locked time courses and spatial activation patterns. This was done for each participant individually. CCA components that showed HFO activity around 20 ms post-stimulus *and* a spatial activation pattern indicating a tangentially oriented dipole were selected for further analysis (referred to as *tangential CCA component*; Fig. 2C). In contrast, CCA components that showed a unipolar activation pattern over the right primary somatosensory cortex were classified as *radial CCA components*. The tangential CCA components were always found to be the CCA components with the highest canonical correlation coefficients, while the radial CCA components were always second. For a more detailed description of the CCA decomposition approach, please refer to our original article (Stephani et al., 2021). Using this approach to obtain spatial filters that optimized for stimulus-locked high-frequency activity, HFO were possible to observe in 27 out of 32 participants (indicated by visual inspection of spatial activation patterns and single-trial, as well as average evoked responses). In order to extract the amplitude of HFO on a single-trial basis, we quantified the area-under-the-curve (AUC) of the Hilbert envelope, measured in a time interval of ±3 ms around the latency of the N20 component (individually determined based on the within-participant average SEP). In an additional analysis, we split that time window and extracted the AUC of the HFO envelope both in the window 3 ms before and after the individually determined N20 peak separately (referred to as pre– and post-N20 HFO).

**Fig. 2.**
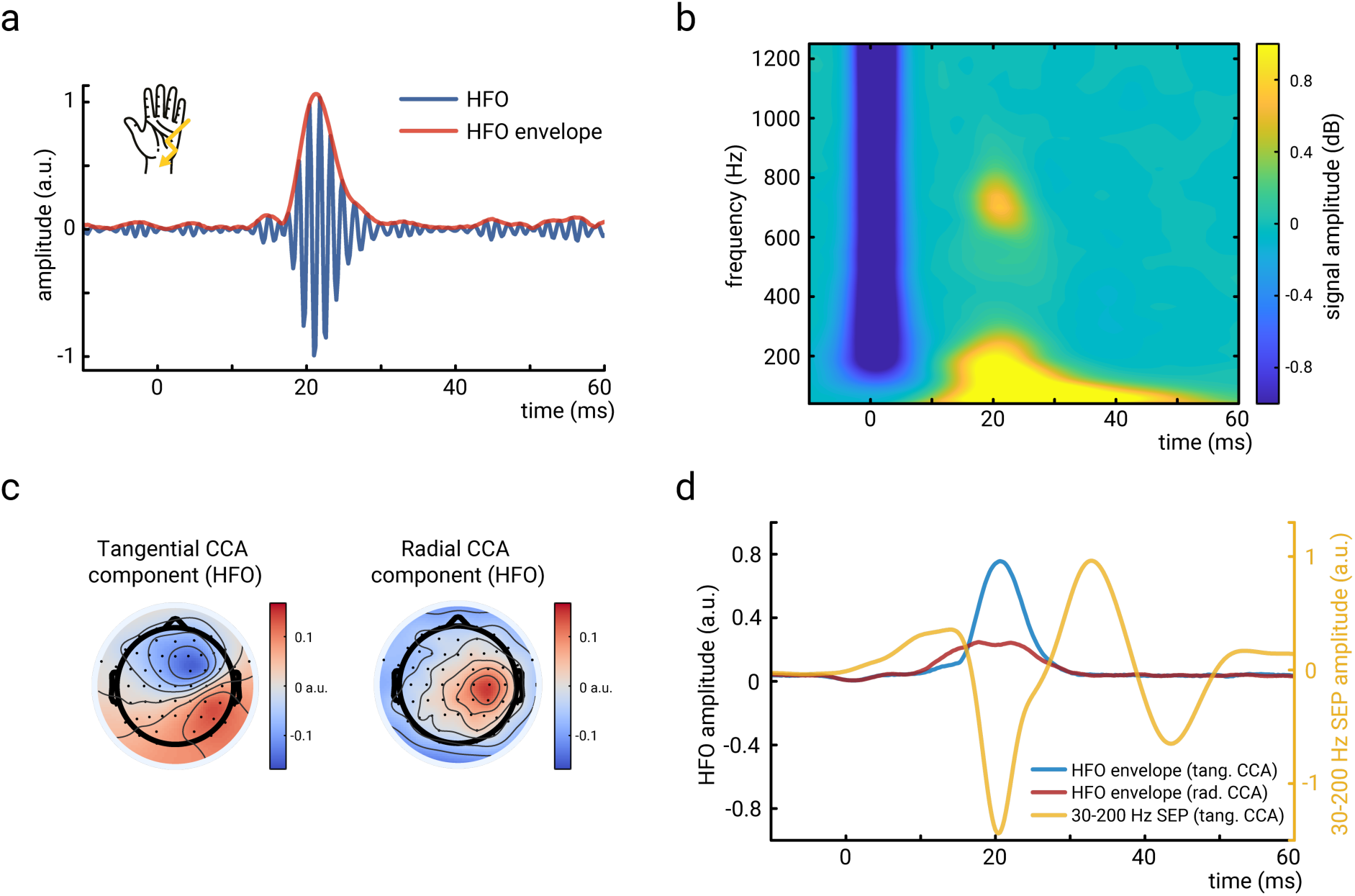
High-frequency oscillations (HFO) extracted with canonical correlation analysis (CCA). a) HFO of an exemplary participant, averaged over 1000 trials. Displayed are both the stimulus-evoked oscillations as well as their signal envelope (extracted using Hilbert transform). Activity derived from the tangential component of our CCA-based spatial filtering approach. b) Time-frequency representation of the short-latency responses of the somatosensory cortex (averaged across participants; N=27). HFO are visible at 20ms in the frequency range around 700 Hz and are clearly distinct from the conventional SEP in lower frequencies. As in panel a), activity derived from the tangential CCA component. c) Spatial activation patterns of the tangential and radial CCA components computed for HFO (averaged across participants; N=27). d) Grand averages of the HFO envelopes (tangential and radial CCA components) as well as the conventional SEP in the frequency range from 30 to 200 Hz (averaged across participants; N=27). Please note that separate CCAs were performed for HFO and the 30-to-200-Hz SEP (for the latter, see Stephani et al., 2021), both resulting in similar tangential and radial components.

Time-frequency representations of HFO were computed using complex Morlet wavelets between 40 and 1250 Hz, with cycle numbers logarithmically scaled from 3 to 20 (60 frequencies in total).

For the visualization and cluster-based permutation tests of the average HFO envelopes of different behavioral response categories, additional sub-sampling of the trials was required since response categories differed in their numbers (arithmetic means of trial numbers per response category: *N*_”strong”|strong_=341.9; *N*_”weak”|strong_=145.6; *N*_”strong”|weak_=140.0; *N*_”weak”|weak_=349.4). As could be expected, higher trial numbers resulted in a lower noise level (and thus lower baseline activity) of the averaged HFO envelopes than lower trial numbers. Trial sub-sampling was performed as follows (individually performed for each participant): The amount of trials in the response condition with the least trials was randomly selected from the other response conditions and averaged. This procedure was repeated 1000 times and the resulting sub-averages were in turn averaged, in order to account for the random selection of trials that went into the sub-averages and to obtain stable estimates of the HFO envelopes in response conditions with different trial numbers. Single-trial statistics were performed without sub-sampling since the factors *presented stimulus intensity* and *perceived stimulus intensity* were explicitly modelled here and the effects of unequal trial numbers should cancel each other out, given the fully crossed experimental design.

### Statistics

Generalized linear mixed-effects models were employed to examine the relationships between *HFO amplitude*, *presented stimulus intensity*, *N20 amplitude*, *pre-stimulus alpha amplitude*, as well as *perceived stimulus intensity* on a single-trial level. For reasons of model convergence and comparability across models, random-intercept models were computed in all cases (with random factor *subject*). We tested the following models:

(1) *HFO* ∼ 1 + *presented stimulus intensity* + (1 | *subject*),
(2) *Perceived stimulus intensity* ∼ 1+ *presented stimulus intensity* + *prestimulus alpha* + *N20* + *HFO* + (1 | *subject*),
(3) *Perceived stimulus intensity* ∼ 1+ *presented stimulus intensity* + *HFO* + (1 | *subject*),
(4) *HFO* ∼ 1 + *presented stimulus intensity* + *prestimulus alpha* + (1 | *subject*),
(5) *N20* ∼ 1 + *presented stimulus intensity* + *HFO* + (1 | *subject*).

A logit link function was used in case of *perceived stimulus intensity* being the dependent variable, in order to account for its dichotomous scale. Before any statistics, pre-stimulus alpha amplitudes were log-transformed in order to approximate a normal distribution, and all continuous variables were z-transformed. Furthermore, to ensure that no longitudinal trends over the course of the experiment drove the effects of interest (time-on-task effect; Benwell et al., 2019), we detrended *prestimulus alpha amplitudes*, *N20 amplitudes*, and *HFO amplitudes* across trials by regressing out the predictor *trial_index* before subjecting these measures to the generalized linear-mixed effects models described above.

Model comparisons were evaluated using the likelihood ratio test (LRT), the Akaike Information Criterion (AIC), and the Bayesian Information Criterion (BIC). The significance level for all analyses was set to *p* = .05 (two-sided).

Complementarily, Bayesian statistics were based on the regression models (3), (4), and (5) described above, and computed with 4 Markov chain Monte Carlo (MCMC) chains of 6000 iterations each. Priors for the fixed effects of interest were chosen to be normally distributed around a mean of 0 with a standard deviation of 0.1 based on the scaling of effects in Stephani et al., 2021 (i.e., corresponding to a weakly informative prior; McElreath, 2020).

Cluster-based permutation tests were performed for additional testing of the relationships between *HFO amplitude* and *presented stimulus intensity* as well as *perceived stimulus intensity*. For the latter, separate analyses were conducted for strong and weak presented stimuli. In all cases, 10000 permutations of the condition labels of interest were conducted, group-level paired-sample *t*-tests were used to compare the resulting condition differences, and clusters were formed with a pre-threshold of *p*_pre_ = .05, thus resulting in the null distribution for the hypothesis test (sum of *t* values in each cluster). The empirical *t*-value sum was subsequently compared with this null distribution and deemed significant if being more extreme than 5% of the values of the null distribution, corresponding to a cluster significance level of *p*_cluster_ = .05. In addition, we performed the same cluster analysis a second time but with *p*_cluster_ = .1 for the relationship between HFO and perceived stimulus intensity, in order to decrease the type II error of not detecting a true difference (in the context of our interpretation of the null effect here). In order to account for different trial numbers of correct and incorrect behavioral responses, we performed a trial sub-sampling approach for the cluster-based permutation tests regarding the effects on perceived stimulus intensity. This was required since we here split the analysis into strong and weak presented stimulus intensities, which resulted in unbalanced trial numbers for weak stimuli being rated as weak stimuli as compared to weak stimuli being rated as strong stimuli (and correspondingly for the strong presented stimuli; note that the task accuracy rate was around 70% in both conditions – thus markedly above chance level – leading to reduced trial numbers of the incorrect responses [conditions *“weak”|strong* and *“strong”|weak*]). We performed the trial sub-sampling for every participant separately in the following way: First, we drew as many trials from all four behavioral conditions (presented intensity x perceived intensity) as the condition with the fewest trials contained. These trials were randomly drawn (with replacement) and the procedure was repeated 1000 times. Subsequently, we averaged the HFO signals within each of these sub-samples and computed their Hilbert envelopes. The arithmetic mean of these 1000 sub-sampling envelopes then served as our participant-level average HFO envelope, which was subjected to the cluster-based permutation statistics. Also, we use these trial-number controlled HFO envelopes for visualization in Figure 4.

The permutation-based analyses were performed in MATLAB (version 2022b, The MathWorks Inc, Natick, MA). The generalized linear mixed-effects models were calculated in R (version 4.2.2, R Development Core Team, 2018). For the frequentist approach, the lme4 package (version 1.1–30, Bates et al., 2015) was used, estimating the fixed-effect coefficients based on the restricted maximum likelihood (REML). To derive a p value for the fixed-effect coefficients, the denominator degrees of freedom were adjusted using Satterthwaite’s method (Satterthwaite, 1946) as implemented in the R package lmerTest (version 3.1–3, Kuznetsova et al., 2017). The Bayesian analyses were performed with the brms package (version 2.17.0, Bürkner, 2017).

### Open code and open data statements

The code for data analysis as well as the data used for summary statistics and for generating the figures are accessible at: https://osf.io/meyza/?view_only=c43d4bc78bac4b72b1674af19c5ddf89. The raw EEG data cannot be made available in a public repository due to the privacy policies for human biometric data according to the European General Data Protection Regulation (GDPR). Further preprocessed data can be obtained from the corresponding author (TS) upon reasonable request and as far as the applicable data privacy regulations allow it.

## Results

### Robust high-frequency oscillations (HFO) around the N20 peak

Participants performed a somatosensory discrimination task, in which they were to distinguish between somatosensory stimuli of two slightly different, supra-threshold intensities (Stephani et al., 2021). Irrespective of the presented and perceived stimulus intensities, clear HFO in response to the median nerve stimuli were observed in 27 out of 32 participants, employing a spatial filtering approach using canonical correlation analysis (CCA; Fig. 2). In all of the 27 participants with overall clear HFO, one CCA component was identified that had a spatial activation pattern suggesting an underlying dipole with tangential orientation – strongly resembling the dipole of the N20 potential (see Stephani et al., 2021) – as well as another CCA component that showed a radial spatial pattern (Fig. 2c). Grand averages of both the tangential and radial CCA components of HFO activity are displayed in Figure 2d. Since our focus here is on HFO that have a cortical origin (Gobbelé et al., 2004) and that are likely generated by the same neuronal populations as the N20 potential (Ikeda et al., 2005), we will continue our analyses with the tangential CCA component in the following sections. Noteworthy, for this tangential CCA component, HFO frequencies were observed to be centered around 700 Hz (Fig. 2b), not 600 Hz as described in previous studies measuring HFO at sensor level above the somatosensory cortex (e.g., Waterstraat et al., 2021).

### HFO relate to presented but not perceived stimulus intensity

The main objective of this study is to examine whether moment-to-moment fluctuations of the perceived stimulus intensity that we observed in our previous study (Stephani et al., 2021) are also reflected in HFO amplitudes.

To this end, we first tested whether HFO were related even to the minute intensity variations of the presented stimuli in our study (i.e., just above the least-noticeable difference) – which should be expected based on previous literature (Klostermann et al., 1998; Gobbelé et al., 2008) yet with the caveat that these studies compared much bigger intensity differences than us. This comparison is required for the further evaluation of the *perceived* stimulus intensity. Our data confirmed that a higher *presented* stimulus intensity was indeed associated with larger HFO amplitudes (Fig. 3a), as tested both with cluster-based permutation statistics (applied to the within-participant averages, *p* < .05; Fig. 3b), as well as with a linear-mixed-effects model operating on single-trial HFO amplitudes, *β* = .031, *t*(26349.0) = 2.534, *p* = .011 (HFO were derived from the tangential CCA component; notably, the size of this intensity effect on HFO was in a similar effect size range as was the case for the N20 component in our previous study, Stephani et al., 2021). As can be seen from Fig. 3b, the presented stimulus intensity modulated HFO amplitudes around their peak (which is congruent with latencies of the N20 peak), with an apparently slightly longer lasting effect after the HFO peak than before their peak.

**Fig. 3.**
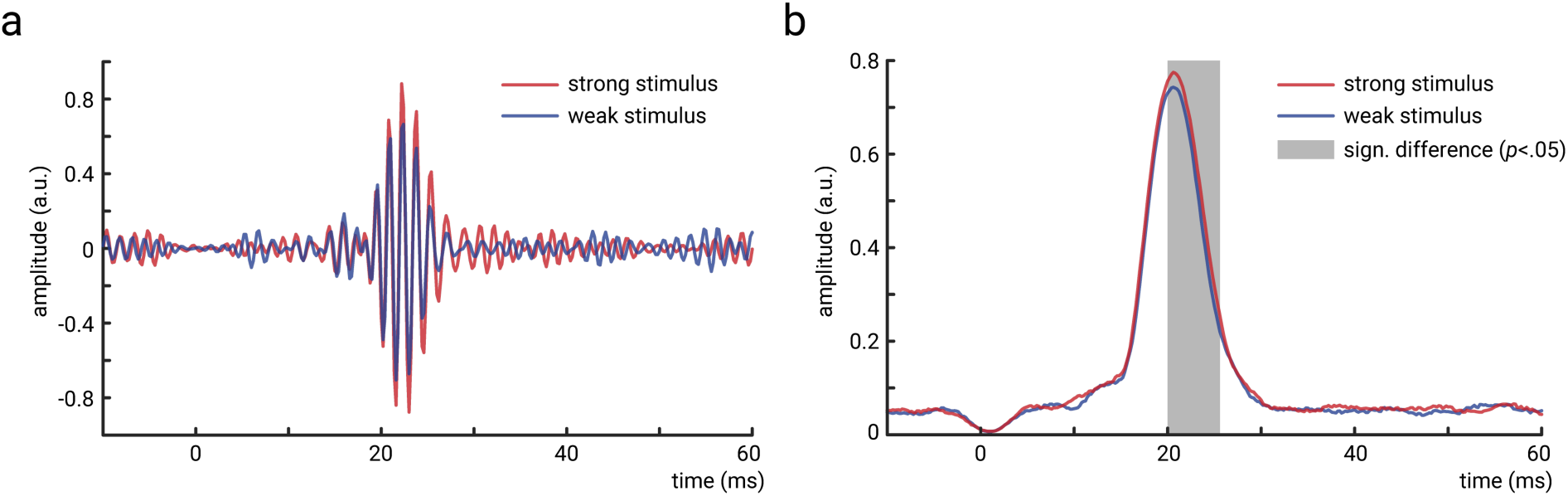
Effect of presented stimulus intensity on HFO amplitudes. a) Single-participant average of HFO for high and low stimulus intensities. b) Group-level average (N=27) of HFO envelopes for high and low stimulus intensities. Permutation-based cluster statistics indicated a difference shortly after ∼20 ms (marked in grey).

Next, we examined the relationship between HFO and perceived stimulus intensity while controlling for the presented stimulus intensity (Fig. 4). For this, we used three approaches: i) adding *HFO amplitude* as additional predictor of *perceived intensity*, next to our previous predictors *pre-stimulus alpha* and *N20 amplitude*, as well as *presented intensity* (Stephani et al., 2021), ii) examining HFO in a separate regression model only including *presented intensity* as a covariate (to account for possible issues of multicollinearity in approach i)), and iii) employing cluster-based permutation statistics separately in the high– and low-intensity conditions.

**Fig. 4.**
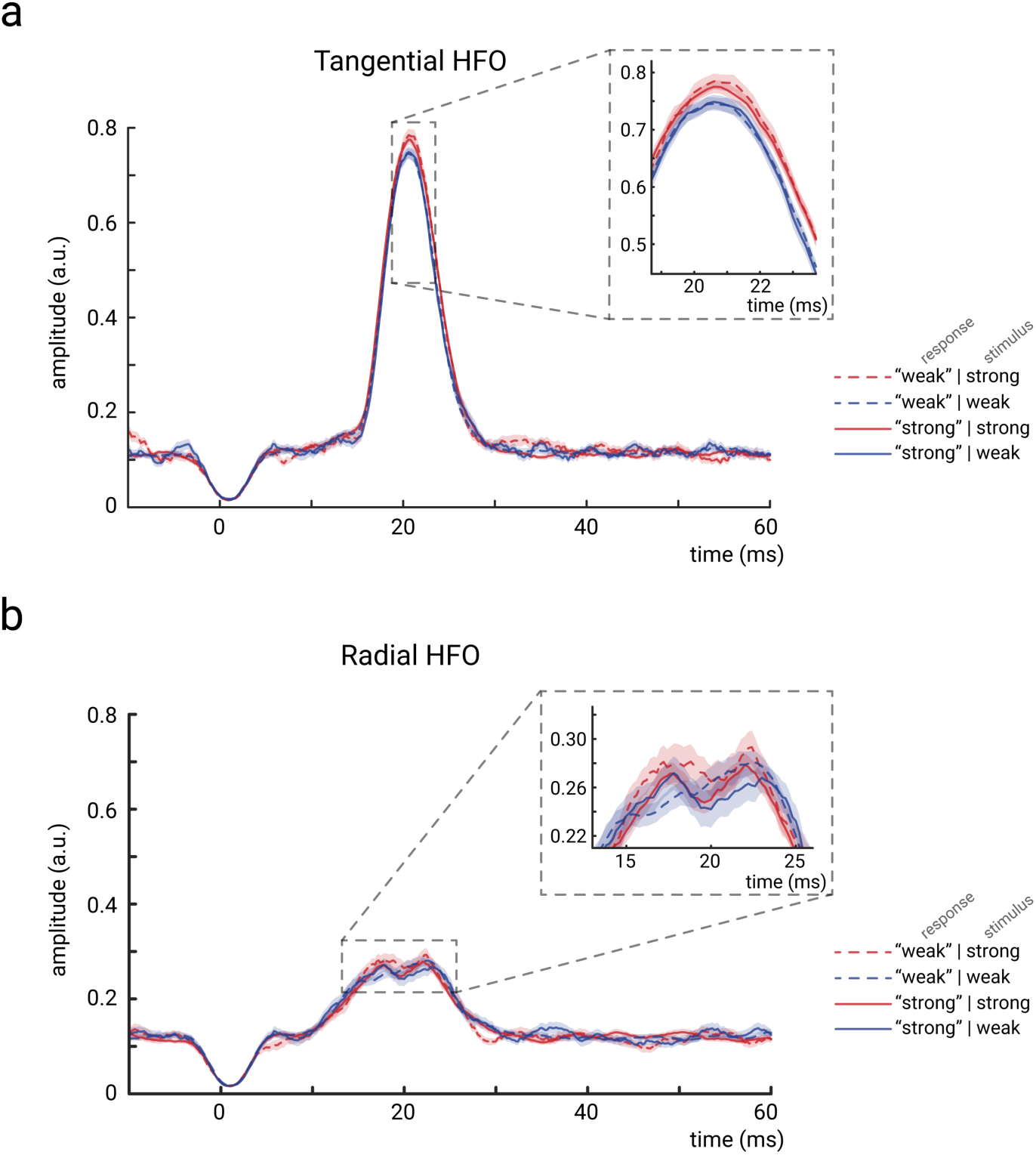
Relationship between HFO and the perceived stimulus intensity. a) No effect of HFO on the *perceived* stimulus intensity can be observed for the tangential CCA component when controlling for the *presented* stimulus intensity. b) Similarly, no effect of HFO on perceived stimulus intensity emerged for the radial CCA component. Please note that trial sub-sampling was performed both in panel a and b, in order to account for different noise levels (i.e., baseline activity) that resulted from smaller trial numbers in the conditions *“weak”|strong* and *“strong”|weak* as compared to *“weak”|weak* and *“strong”|strong*. All curves reflect group-level averages of HFO envelopes; shaded areas correspond to the standard error of the mean based on the within-subject variances (Morey, 2008).

Adding HFO to our previous regression model did not lead to a better model fit, thus not suggesting any relevance of HFO for perceived stimulus intensity beyond the previously identified predictors, *χ*^2^(1) = 0.793, *p* = .373, AIC_incl. HFO_ = 31170 vs. AIC_excl. HFO_ = 31169, BIC_incl. HFO_ = 31219 vs. BIC_excl. HFO_ = 31210. Furthermore, HFO effects were not significant in the dedicated regression models only considering HFO and presented stimulus intensity as predictors, *β* = .012, *z* = 0.860, *p* = .390. We evaluated these non-significant effects with additional Bayesian statistics, which indicated moderate evidence for the absence of a relationship between HFO and perceived stimulus intensity, BF01 = 4.71, *β*_Bayes_ = .013, 95%-credibility interval = [-.014, .040]. Similar evidence was provided by the cluster-based permutation statistics, which did not show any effect of HFO on perceived stimulus intensity either in the high-or in the low-presented-intensity condition (even with the liberal threshold of a cluster significance level of *p* < .1).

Taken together, spontaneous fluctuations of HFO amplitude thus do not seem to be related to any changes of the *perceived* intensity in our somatosensory discrimination paradigm (when controlling for the physically presented intensity).

### HFO are not affected by pre-stimulus alpha amplitude and only weakly related to N20 peaks

Since previous work found relationships between pre-stimulus alpha band amplitude and stimulus-evoked activity in conventional frequency bands, we tested for such a relationship also in HFO activity. The same pre-stimulus alpha activity that affected N20 amplitudes in our previous study (Stephani et al., 2021) did not show any effect on HFO amplitude, as tested with a linear-mixed effects model, *β* = –.007, *t*(24106.4) = –0.957, *p* = .338. Furthermore, Bayesian statistics provided moderate to strong evidence for the absence of an effect, BF01 = 8.65, *β*_Bayes_ = –.007, 95%-credibility interval = [-.021, .007]. (Very similar statistical results were obtained from the LMEs when not using the pre-stimulus alpha metric from our previous study but instead pre-stimulus alpha activity derived from the CCA filters trained to detect HFO for the current study, *β* = –.007, *t*(20834.2) = –0.900, *p* = .368.)

HFO and N20 amplitudes did show a significant relationship, *β* = –.013, *t*(26352.7) = –2.457, *p* = .014 (frequentist LME analysis), yet corresponding to only weak evidence for H1 according to the Bayesian approach, BF10 = 1.14, *β*_Bayes_ = –.013, 95%-credibility interval = [-.022, –.002]. The negative effect direction indicates that larger HFO amplitudes were associated with more negative N20 peak amplitudes (i.e. larger magnitudes) which is also the effect direction that should be expected based on the effects of presented stimulus intensity on both N20 and HFO (but please note that we included stimulus intensity as a covariate in this analysis; thus, the weak yet significant relationship between N20 and HFO remains beyond trivial effects of stimulus intensity changes).

In summary, HFO amplitudes did thus not covary at all with the previously identified marker of excitability fluctuations in alpha amplitude fluctuations and were only weakly associated with the N20 peak amplitude of the conventional event-related response, which emerges at the very same time but in a lower frequency band.

### Comparison of pre-versus post-N20 HFO

In order to exclude the possibility that our analyses of tangential HFO did not only reflect intra-cortical processes of somatosensory pyramidal cells but also pre-synaptic signals originating from thalamocortical axon terminals with a similar spatial orientation (Gobbelé et al., 2004), we additionally examined HFO occurring in the 3ms *after* the N20 peak only. Results of the Bayesian linear-mixed-effects models for the relationships between HFO and perceived stimulus intensity, HFO and N20, as well as HFO and pre-stimulus alpha amplitude yielded very similar results as in above analyses for the whole time window, BF01_perceived_ = 6.28, *β*_perceived_ = .008, 95%-credibility interval_perceived_ = [-.019, .035], BF01_N20_ = 2.61, *β*_N20_ = –.011, 95%-credibility interval_N20_ = [-.021, .000], as well as BF01_prestim_ = 7.75, *β*_prestim_ = –.008, 95%-credibility interval_prestim_ = [-.022, .006]. When testing pre-N20 HFO (in a time window starting 3 ms before the N20 peak), no other effects emerged either, with Bayes factors being similar or even higher in favor of the null hypotheses, BF01_perceived_ = 5.26, BF01_N20_ = 5.95, BF01_prestim_ = 10.6. Thus, it is unlikely that the lack of relation between HFO and our other measures of interest was due to a mixed nature of pre– and post-synaptic HFO. Furthermore, this post-hoc analysis suggests that our findings are valid even when isolating the late part of HFO that can only be of intra-cortical origin, that is, presumably free from any thalamic contribution.

## Discussion

Re-examining an EEG dataset of a somatosensory intensity discrimination paradigm (Stephani et al., 2021), we here tested the involvement of neuronal population spikes, as measured by high-frequency oscillations (HFO), in moment-to-moment dynamics of neural excitability and their effects on perceived stimulus intensity. Using Bayesian statistics, we found evidence against the involvement of HFO in behaviorally relevant excitability dynamics. Thus, population spiking activity of first feedforward cortical processing cannot account for state-dependent modulation of sensory processes. Instead, following the notion that conventional evoked potentials in lower frequencies – such as the N20 component (Stephani et al., 2021) – may reflect a backpropagation of membrane potentials towards the apical dendrites, we suggest that top-down modulatory processes (e.g., related to alpha oscillations) take place at distal sites of the involved pyramidal cells rather than at basal compartments.

### Evidence against the involvement of population spiking activity in the modulation of perceived stimulus intensity

As expected and in line with previous work on HFO in humans (Klostermann et al., 1998; Gobbelé et al., 2008), we observed larger HFO amplitudes for higher *presented* stimulus intensities. However, in contrast to previous studies, the presented stimulus intensities used here differed only very slightly regarding their somatosensory percept since we deliberately adjusted them to be close to the *just-noticeable difference*. Thus, even minute differences in the presented stimulus intensity are reflected in HFO amplitudes, which validates the sensitivity of our non-invasive measure of population spiking activity. However, despite this robust relationship between HFO and presented stimulus intensity, we did not observe any association of trial-to-trial fluctuations of HFO and the variability of the *perceived* stimulus intensity (i.e., when controlling for the presented stimulus intensity), neither with generalized linear-mixed effects modeling, nor cluster-based permutation tests. Moreover, Bayesian statistics provided even moderate evidence *against* the involvement of HFO in the modulation of perceived stimulus intensity. We also tested this for post-N20 HFO separately, thus making sure that our findings cannot be biased by pre-synaptic HFO from thalamocortical axon terminals (Gobbelé et al., 2004; Ritter et al., 2008), leading to the same result of perceptual variability at the time of the N20 yet without involvement of HFO.

### Cortical task-related HFO in humans reside at frequencies around 700 Hz

Conventionally, HFO in the somatosensory domain of humans are assumed to arise in a frequency band around 600 Hz, so-called σ bursts (Curio, 2000; Waterstraat et al., 2016). Our data in contrast suggested slightly higher frequencies, centered at 700 Hz (Fig. 2b). At first, this may seem at odds, yet previous studies which also focused on tangentially oriented σ bursts in fact reported HFO peak frequencies clearly higher than 600 Hz and rather approaching 700 Hz (e.g., Hashimoto et al., 1996; Waterstraat et al., 2016; Waterstraat et al., 2021). Additionally, when studying a presumably mixed signal of both tangential and radial HFO, peak frequencies also exceeded 600 Hz (Ritter et al., 2008). In the latter study, an increase of HFO frequency was associated with a decrease in stimulation frequency, emphasizing the role of the inter-stimulus interval as has also been observed by Klostermann et al. (1999). Given that our stimulation frequency was fairly low (on average 0.66 Hz), which should preclude strong trial-to-trial attenuation effects, we conclude that in situations where somatosensory stimuli are perceived in isolation (and not in the form of a pulse train as in passive stimulation paradigms; typically using repetition rates of around 3 Hz, e.g. Waterstraat et al., 2021), cortical HFO may in fact reside in a higher frequency range than the conventionally reported 600 Hz. This may thus be an important observation to guide future studies that examine HFO in psychophysics paradigms that require behavioral responses.

### Excitability-related modulation of sensory input may emerge through backpropagating membrane potentials

If neuronal population spiking activity is not involved in the behaviorally relevant variability of initial cortical processing found in previous work (Stephani et al., 2021), which are the neuronal processes that mediate these effects? Following the notion that HFO are generated at basal regions of cortical pyramidal cells (Shimazu et al., 2000; Curio, 2000; Baker et al., 2003; Hu et al., 2023) while the N20 component reflects post-synaptic membrane potentials that propagate “back” in soma-dendritic direction (Wikström et al., 1996; Bruyns-Haylett et al., 2017; Thorpe et al., 2024), our data are in line with the interpretation that spontaneous neuronal states shape intraneuronal depolarization along the apical dendrites, instead of the output firing activity at basal pyramidal cell compartments (HFO) – at least not during the initial cortical feedforward volley. This may also explain why the relation between alpha band oscillations, a commonly used marker for cortical excitability states, and neuronal firing activity are typically found to be rather small, even in animal work with invasive recordings (e.g., Haegens et al., 2011). Crucially, this interpretation is based on the assumption that the intracortical component of HFO originates from the same pyramidal cells as the N20 potential, as has previously been suggested using local field potentials, extracellular single unit recordings and surface EEG in non-human animals (Shimazu et al., 2000; Baker et al., 2003; Ikeda et al., 2005). Furthermore, HFO are not an artifact of electrical nerve stimulation but are present in naturalistic tactile stimulation as well (Baker et al., 2003).

Although it may seem puzzling that behaviorally relevant variability of low-frequency SEP activity, which reflects post-synaptic membrane potentials (Lopes da Silva, 2004; Ilmoniemi and Sarvas, 2019; Næss et al., 2021), does not directly relate to the neurońs firing output, our observations of only weakly linked N20 and HFO dynamics complement the previous literature very well. For instance, it has been reported that the N20 shows small increments when entering deep sleep stages while HFO are drastically diminished (Hashimoto et al., 1996). Furthermore, the N20 and HFO are affected differentially by propofol anesthesia: Whereas the N20 remains stable, HFO are substantially decreased both in amplitude and frequency (Klostermann et al., 2000). Additionally, HFO were found to be increased in eyes-open as compared to eyes-closed EEG recordings while the N20 appears to be unaffected (Gobbelé et al., 2008). In contrast, isometric motor contraction leads to an attenuation of the N20 but not of HFO (Klostermann et al., 2001). Such motor gating of the somatosensory domain has been proposed to emerge via efference copies (Holst, 1954; Wolpert and Ghahramani, 2000; Palmer et al., 2016; Kilteni et al., 2020; Job and Kilteni, 2023) and may act via layer 1 top-down projections (Manita et al., 2015; Manita et al., 2017), further supporting the notion of functionally distinct dendritic compartments of cortical pyramidal cells. Last but not least, transcranial alternating current stimulation (tACS) at alpha frequencies has been reported to up-regulate N20 amplitudes but not HFO (Fabbrini et al., 2022). Together, these studies provide compelling evidence of the differential functional properties of the N20 component and HFO.

Moreover, the notion of a neuronal state-dependent modulation of backpropagating membrane potentials in apical dendrites towards the superficial cortical layers is not only well in line with Dendritic Integration Theory which hypothesizes the interaction between apical and basal dendritic compartments to be pivotal for conscious stimulus perception (Aru et al., 2020; Bachmann et al., 2020; Larkum, 2022), but also complies with recent observations of alpha band activity exerting a top-down influence on superficial layers of primary sensory regions (Halgren et al., 2019), as well as with earlier conceptions of the interplay between re-entrant bottom-up and top-down signals in superficial layers at early stages of the somatosensory response cascade (Cauller and Kulics, 1991; Cauller, 1995). In this context, it is noteworthy that also HFO result in membrane potentials that travel in soma-dendritic direction (Ikeda et al., 2005), yet at the frequency of HFO and thus possibly resembling back-propagating action potentials (Buzsáki and Kandel, 1998; Waters et al., 2005). Although it is conceivable that our HFO signals also reflected these additional, indirect manifestations of the neuronal output, this does not weaken our interpretation since the back-propagating action potentials are expected to be proportional to the outgoing action potentials.

It remains an open question, however, what happens to the backpropagating post-synaptic potentials that are visible in the EEG as the N20 component. Which subsequent neuronal processes use this information and in what way does this affect sensory information processing down-stream? A tentative hypothesis may be that the backpropagated N20 potential serves the function of a “neuronal buffer”, that is, a trace of the immediate past of sensory exposure within specific pyramidal cells/ pyramidal cell populations. In line with previously observed trial-to-trial temporal dependencies for perceptual decision making in general (Urai et al., 2019; Forster et al., 2024) and for early somatosensory potentials in particular (Stephani et al., 2020; Stephani et al., 2022), this information may then be integrated with both top-down as well as subsequent bottom-up sensory input. Yet, to test this, approaches are required to selectively modulate apical dendritic activity in humans non-invasively.

## Acknowledgements

We very much thank Gabriel Curio and Gunnar Waterstraat for discussions of the HFO results presented in this manuscript.

## References

1. Allison T, McCarthy G, Wood CC, Jones SJ (1991) Potentials Evoked in Human and Monkey Cerebral Cortex by Stimulation of the Median Nerve. Brain 114:2465–2503.

2. Arieli A, Sterkin A, Grinvald A, Aertsen A (1996) Dynamics of ongoing activity. Explanation of the large variability in evoked cortical responses. Science 273:1868–1871.

3. Aru J, Suzuki M, Larkum ME (2020) Cellular Mechanisms of Conscious Processing. Trends Cogn Sci 24:814–825.

4. Bachmann T, Suzuki M, Aru J (2020) Dendritic integration theory: A thalamo-cortical theory of state and content of consciousness. PhiMiSci 1:2.

5. Baker SN, Curio G, Lemon RN (2003) EEG oscillations at 600 Hz are macroscopic markers for cortical spike bursts. J Physiol (Lond) 550:529–534.

6. Bates D, Mächler M, Bolker B, Walker S (2015) Fitting Linear Mixed-Effects Models Using lme4. J. Stat. Soft. 67:1–48.

7. Benwell CS, London RE, Tagliabue CF, Veniero D, Gross J, Keitel C, Thut G (2019) Frequency and power of human alpha oscillations drift systematically with time-on-task. Neuroimage 192:101– 114.

8. Bruyns-Haylett M, Luo J, Kennerley AJ, Harris S, Boorman L, Milne E, Vautrelle N, Hayashi Y, Whalley BJ, Jones M, Berwick J, Riera J, Zheng Y (2017) The neurogenesis of P1 and N1. A concurrent EEG/LFP study. Neuroimage 146:575–588.

9. Bürkner P-C (2017) brms: An R Package for Bayesian Multilevel Models Using Stan. J. Stat. Soft. 80:1–28.

10. Buzsáki G, Kandel A (1998) Somadendritic backpropagation of action potentials in cortical pyramidal cells of the awake rat. J Neurophysiol 79:1587–1591.

11. Cauller L (1995) Layer I of primary sensory neocortex: where top-down converges upon bottom-up. Behav Brain Res 71:163–170.

12. Cauller LJ, Kulics AT (1991) The neural basis of the behaviorally relevant N1 component of the somatosensory-evoked potential in SI cortex of awake monkeys: evidence that backward cortical projections signal conscious touch sensation. Exp Brain Res 84:607–619.

13. Curio G (2000) Linking 600-Hz “spikelike” EEG/MEG wavelets (“sigma-bursts”) to cellular substrates: concepts and caveats. Journal of Clinical Neurophysiology 17:377–396.

14. Delorme A, Makeig S (2004) EEGLAB. An open source toolbox for analysis of single-trial EEG dynamics including independent component analysis. J Neurosci Methods 134:9–21.

15. Fabbrini A, Guerra A, Giangrosso M, Manzo N, Leodori G, Pasqualetti P, Conte A, Di Lazzaro V, Berardelli A (2022) Transcranial alternating current stimulation modulates cortical processing of somatosensory information in a frequency– and time-specific manner. Neuroimage 254:119119.

16. Fedele T, Scheer H-J, Burghoff M, Waterstraat G, Nikulin VV, Curio G (2013) Distinction between added-energy and phase-resetting mechanisms in non-invasively detected somatosensory evoked responses. Conference proceedings: IEEE Engineering in Medicine and Biology Society. Annual Conference 2013:1688–1691.

17. Forster C, Stephani T, Grund M, Panagoulas E, Al E, Hofmann SM, Nikulin VV, Villringer A (2024) Pre-stimulus beta power encodes explicit and implicit perceptual biases in distinct cortical areas. bioRxiv. doi:10.1101/2024.06.12.598458.

18. Gobbelé R, Dieckhöfer A, Thyerlei D, Buchner H, Waberski TD (2008) The impact of stimulus properties on low– and high-frequency median nerve somatosensory evoked potentials. J Clin Neurophysiol 25:194–201.

19. Gobbelé R, Waberski TD, Simon H, Peters E, Klostermann F, Curio G, Buchner H (2004) Different origins of low– and high-frequency components (600 Hz) of human somatosensory evoked potentials. Clinical Neurophysiology 115:927–937.

20. Green DM, Swets JA (1966) Signal detection theory and psychophysics. New York, NY: Wiley.

21. Haegens S, Nácher V, Luna R, Romo R, Jensen O (2011) α-Oscillations in the monkey sensorimotor network influence discrimination performance by rhythmical inhibition of neuronal spiking. Proc Natl Acad Sci U S A 108:19377–19382.

22. Halgren M, Ulbert I, Bastuji H, Fabó D, Erőss L, Rey M, Devinsky O, Doyle WK, Mak-McCully R, Halgren E, Wittner L, Chauvel P, Heit G, Eskandar E, Mandell A, Cash SS (2019) The generation and propagation of the human alpha rhythm. Proc Natl Acad Sci U S A 116:23772–23782.

23. Hashimoto I, Mashiko T, Imada T (1996) Somatic evoked high-frequency magnetic oscillations reflect activity of inhibitory interneurons in the human somatosensory cortex. Electroencephalography and Clinical Neurophysiology/Evoked Potentials Section 100:189–203.

24. Haufe S, Meinecke F, Görgen K, Dähne S, Haynes J-D, Blankertz B, Bießmann F (2014) On the interpretation of weight vectors of linear models in multivariate neuroimaging. Neuroimage 87:96–110.

25. Holst E von (1954) Relations between the central Nervous System and the peripheral organs. The British Journal of Animal Behaviour 2:89–94.

26. Hu H, Hostetler RE, Agmon A (2023) Ultrafast (400 Hz) network oscillations induced in mouse barrel cortex by optogenetic activation of thalamocortical axons. Elife 12:e82412.

27. Ikeda H, Wang Y, Okada YC (2005) Origins of the somatic N20 and high-frequency oscillations evoked by trigeminal stimulation in the piglets. Clin Neurophysiol 116:827–841.

28. Ilmoniemi R, Sarvas J (2019) Brain signals. Physics and mathematics of MEG and EEG. Cambridge: The MIT Press.

29. Job X, Kilteni K (2023) Action does not enhance but attenuates predicted touch. Elife 12.

30. Kilteni K, Engeler P, Ehrsson HH (2020) Efference Copy Is Necessary for the Attenuation of Self-Generated Touch. iScience 23:100843.

31. Klostermann F, Funk T, Vesper J, Siedenberg R, Curio G (2000) Propofol narcosis dissociates human intrathalamic and cortical high-frequency ( 400 hz) SEP components. Neuroreport 11:2607–2610.

32. Klostermann F, Gobbele R, Buchner H, Siedenberg R, Curio G (2001) Differential gating of slow postsynaptic and high-frequency spike-like components in human somatosensory evoked potentials under isometric motor interference. Brain Research 922:95–103.

33. Klostermann F, Nolte G, Curio G (1999) Multiple generators of 600 Hz wavelets in human SEP unmasked by varying stimulus rates. Neuroreport 10:1625–1629.

34. Klostermann F, Nolte G, Losch F, Curio G (1998) Differential recruitment of high frequency wavelets (600 Hz) and primary cortical response (N20) in human median nerve somatosensory evoked potentials. Neurosci Lett 256:101–104.

35. Kulics AT, Cauller LJ (1986) Cerebral cortical somatosensory evoked responses, multiple unit activity and current source-densities: their interrelationships and significance to somatic sensation as revealed by stimulation of the awake monkey’s hand. Exp Brain Res 62:46–60.

36. Kuznetsova A, Brockhoff PB, Christensen RHB (2017) lmerTest Package: Tests in Linear Mixed Effects Models. J. Stat. Soft. 82:1–26.

37. Larkum M (2013) A cellular mechanism for cortical associations: an organizing principle for the cerebral cortex. Trends in Neurosci 36:141–151.

38. Larkum M (2022) Are dendrites Conceptually Useful? Neuroscience 489:4–14.

39. Larkum ME, Kaiser KM, Sakmann B (1999) Calcium electrogenesis in distal apical dendrites of layer 5 pyramidal cells at a critical frequency of back-propagating action potentials. Proc Natl Acad Sci U S A 96:14600–14604.

40. Larkum ME, Senn W, Lüscher H-R (2004) Top-down dendritic input increases the gain of layer 5 pyramidal neurons. Cereb Cortex 14:1059–1070.

41. Lopes da Silva F (2004) Functional localization of brain sources using EEG and/or MEG data: volume conductor and source models. Magn Reson Imaging 22:1533–1538.

42. Manita S, Miyakawa H, Kitamura K, Murayama M (2017) Dendritic Spikes in Sensory Perception. Front Cell Neurosci 11:29.

43. Manita S, Suzuki T, Homma C, Matsumoto T, Odagawa M, Yamada K, Ota K, Matsubara C, Inutsuka A, Sato M, Ohkura M, Yamanaka A, Yanagawa Y, Nakai J, Hayashi Y, Larkum ME, Murayama M (2015) A Top-Down Cortical Circuit for Accurate Sensory Perception. Neuron 86:1304–1316.

44. Marvan T, Polák M, Bachmann T, Phillips WA (2021) Apical amplification—a cellular mechanism of conscious perception? Neurosci Conscious 2021:niab036.

45. McCormick DA, Nestvogel DB, He BJ (2020) Neuromodulation of Brain State and Behavior. Annu Rev Neurosci 43:391–415.

46. McElreath R (2020) Statistical rethinking. A Bayesian course with examples in R and Stan. Boca Raton, London, New York: CRC Press.

47. Morey RD (2008) Confidence Intervals from Normalized Data: A correction to Cousineau (2005). Tutorial in Quantitative Methods for Psychology 4:61–64.

48. Næss S, Halnes G, Hagen E, Hagler DJ, Dale AM, Einevoll GT, Ness TV (2021) Biophysically detailed forward modeling of the neural origin of EEG and MEG signals. Neuroimage 225:117467.

49. Nicholson Peterson N, Schroeder CE, Arezzo JC (1995) Neural generators of early cortical somatosensory evoked potentials in the awake monkey. Electroencephalography and Clinical Neurophysiology/Evoked Potentials Section 96:248–260.

50. Oldfield RC (1971) The assessment and analysis of handedness: The Edinburgh inventory. Neuropsychologia 9:97–113.

51. Ozaki I, Hashimoto I (2011) Exploring the physiology and function of high-frequency oscillations (HFOs) from the somatosensory cortex. Clin Neurophysiol 122:1908–1923.

52. Palmer CE, Davare M, Kilner JM (2016) Physiological and Perceptual Sensory Attenuation Have Different Underlying Neurophysiological Correlates. J Neurosci 36:10803–10812.

53. Phillips WA, Clark A, Silverstein SM (2015) On the functions, mechanisms, and malfunctions of intracortical contextual modulation. Neurosci Biobehav Rev 52:1–20.

54. Ritter P, Freyer F, Curio G, Villringer A (2008) High-frequency (600 Hz) population spikes in human EEG delineate thalamic and cortical fMRI activation sites. Neuroimage 42:483–490.

55. Sadaghiani S, Hesselmann G, Friston KJ, Kleinschmidt A (2010) The relation of ongoing brain activity, evoked neural responses, and cognition. Front Syst Neurosci 4:20.

56. Samaha J, Iemi L, Haegens S, Busch NA (2020) Spontaneous Brain Oscillations and Perceptual Decision-Making. Trends Cogn Sci 24:639–653.

57. Satterthwaite FE (1946) An Approximate Distribution of Estimates of Variance Components. Biometrics Bulletin 2:110.

58. Shimazu H, Kaji R, Tsujimoto T, Kohara N, Ikeda A, Kimura J, Shibasaki H (2000) High-frequency SEP components generated in the somatosensory cortex of the monkey. Neuroreport 11.

59. Stephani T, Hodapp A, Jamshidi Idaji M, Villringer A, Nikulin VV (2021) Neural excitability and sensory input determine intensity perception with opposing directions in initial cortical responses. Elife 10:e67838.

60. Stephani T, Nierula B, Villringer A, Eippert F, Nikulin VV (2022) Cortical response variability is driven by local excitability changes with somatotopic organization. Neuroimage 264:119687.

61. Stephani T, Waterstraat G, Haufe S, Curio G, Villringer A, Nikulin VV (2020) Temporal Signatures of Criticality in Human Cortical Excitability as Probed by Early Somatosensory Responses. J Neurosci 40:6572–6583.

62. Thorpe RV, Black CJ, Borton DA, Hu L, Saab CY, Jones SR (2024) Distinct neocortical mechanisms underlie human SI responses to median nerve and laser-evoked peripheral activation. Imaging Neuroscience 2:1–29.

63. Urai AE, Gee JW de, Tsetsos K, Donner TH (2019) Choice history biases subsequent evidence accumulation. Elife 8:e46331.

64. Waters J, Schaefer A, Sakmann B (2005) Backpropagating action potentials in neurones: measurement, mechanisms and potential functions. Prog Biophys Mol Biol 87:145–170.

65. Waterstraat G, Fedele T, Burghoff M, Scheer H-J, Curio G (2015) Recording human cortical population spikes non-invasively--An EEG tutorial. J Neurosci Methods 250:74–84.

66. Waterstraat G, Körber R, Storm J-H, Curio G (2021) Noninvasive neuromagnetic single-trial analysis of human neocortical population spikes. Proc Natl Acad Sci U S A 118:e2017401118.

67. Waterstraat G, Scheuermann M, Curio G (2016) Non-invasive single-trial detection of variable population spike responses in human somatosensory evoked potentials. Clin Neurophysiol 127:1872–1878.

68. Wikström H, Huttunen J, Korvenoja A, Virtanen J, Salonen O, Aronen H, Ilmoniemi RJ (1996) Effects of interstimulus interval on somatosensory evoked magnetic fields (SEFs). A hypothesis concerning SEF generation at the primary sensorimotor cortex. Electroencephalography and Clinical Neurophysiology/Evoked Potentials Section 100:479–487.

69. Wolpert DM, Ghahramani Z (2000) Computational principles of movement neuroscience. Nat Neurosci 3 Suppl:1212–1217.

